# Inference and visualization of multi-scale cell tree for decoding functional diversity with scMustree

**DOI:** 10.1101/2025.04.22.649953

**Authors:** Yunpei Xu, Shaokai Wang, Liqing Ding, Hong-Dong Li, Jianxin Wang

**Author notes:** To whom correspondence should be addressed: Jianxin Wang.

## Abstract

Deciphering cellular heterogeneity and functional diversity is crucial to understanding biological systems and disease mechanisms. Although single-cell RNA sequencing (scRNA-seq) has revolutionized this field, existing methods often fail to characterize the functional associations between cell populations due to the discrete nature of clustering results. We introduce scMustree, a tree-construction algorithm that enables in-depth exploration of functional associations and transitional states among cellular populations. scMustree combines top-down iterative decomposition for high-purity leaf nodes with bottom-up merging based on quantitative functional distances, preserving local continuity and global structural clarity. Benchmarking on real datasets shows that scMustree not only outperforms existing methods in clustering accuracy but also reveals functionally coherent subtypes, anatomically related cell types, and developmentally connected transitions. In case studies, it identifies Alzheimer’s disease-associated microglial subtypes and captures spatial cell distribution patterns of cell types in developed embryos, demonstrating its utility as a powerful tool for multi-scale exploration of cellular heterogeneity.

## Introduction

Single-cell RNA sequencing (scRNA-seq) has revolutionized our ability to resolve cellular heterogeneity within tissue microenvironments^1^. Clustering algorithms, as a cornerstone of scRNA-seq analysis^2^, group cells by transcriptional similarity to enable critical downstream applications, including cell type annotation^3^, differential expression analysis^4^, and intercellular communication inference^5^.

Current clustering approaches fall into two paradigms: Traditional frameworks (e.g., Seurat^6^, Scanpy^7^) remain widely adopted, employing highly variable gene selection with linear dimensionality reduction followed by graph-based community detection. In recent years, building upon these foundations, deep learning-enhanced methods have emerged to further improve clustering performance by capturing complex, nonlinear patterns in the data. For instance, DESC^8^ optimizes feature representation through autoencoders, scDCC^9^ integrates Gaussian mixture models for joint feature learning and clustering, and scGNN^10^ employs graph neural networks to model cellular topological relationships. These methods enhance clustering precision through nonlinear feature extraction.

Despite these advancements, three fundamental challenges persist in single-cell clustering analysis. First, algorithmic dependence on predefined cluster numbers contradicts the biological reality of unknown cell-type quantities^11^. In many biological contexts, the true number of cell types or states is inherently uncertain and may vary across tissues or conditions. Relying on a fixed cluster number not only risks oversimplifying the underlying cellular heterogeneity but may also obscure the discovery of rare or transitional populations. Moreover, some methods determine convergence based on quantifiable optimization objectives, which do not necessarily align with biologically meaningful distinctions, potentially leading to biologically irrelevant groupings.

Second, current methods primarily produce discrete cell partitions rather than revealing intercellular relationships^12^. While such hard clustering effectively groups similar cells, it often overlooks the continuous nature of cellular differentiation and functional transitions. As a result, important biological insights, such as functional similarities^13,14^, lineage trajectories^15,16^, hierarchical structures^17–19^, or subtle gradations between cell states^20,21^, may remain hidden. This limitation underscores the need for approaches capable of capturing both discrete groupings and the underlying relational structure among cells.

Third, evaluation metrics exhibit inherent biases due to resolution discrepancies in reference annotations^22^. Specifically, coarse-grained labels may inadequately assess performance, while fine-grained annotations face challenges from increasing data scales and cellular diversity. Moreover, substantial variations in cellular abundance across distinct types result in a pronounced class imbalance^23^. This biological complexity highlights the critical necessity for tree modeling frameworks capable of integrating multi-scale and multi-resolution analyses to comprehensively interpret single-cell data.

Recent advancements in leveraging tree structures for optimizing single-cell analysis have shown promise. For instance, scSHC^24^ improves hierarchical clustering through statistical significance analysis but critically depends on initial tree quality. Similarly, CHOIR^25^ adopts a top-down hierarchical tree construction approach, integrating random forest classifiers with permutation testing to mitigate over-clustering. However, the top-down initialization process may be influenced by the initial clustering performance and may obscure the inter-cluster relationships at the same hierarchical level. SEAT^26^ establishes a tree based on the cell graph and structural entropy reduction, yet faces fundamental limitations in quantifying intercellular relationships within high-dimensional sparse datasets. CHETAH^27^ and MRtree^28^ require pre-existing reference datasets or multi-resolution clustering results. While Wu et al.^29^ proposed hierarchy-sensitive metrics (wRI/wNMI), their work neither addresses tree construction nor accounts for multi-resolution label impacts.

To address these challenges, we propose scMustree (**s**ingle-**c**ell **Mu**lti-**s**cale **t**ree), a tree-construction algorithm designed to infer and visualize hierarchical structures among cellular populations. Rather than treating cell populations as isolated entities, scMustree organizes them into a multi-scale tree structure, where each node or branch captures distinct functional characteristics or transitional potential. By analyzing the topology of this tree, such as branch lengths, divergence points, and nested substructures, researchers can perform an in-depth exploration of functional associations, lineage progressions, and intermediate cellular states.

Through comprehensive benchmarking across 11 single-cell datasets, scMustree demonstrates superior performance over existing methods in both algorithmic accuracy and biological interpretability. Notably, it identifies previously unannotated cellular subtypes and states, such as lipid-associated microglial states in Alzheimer’s disease and activated muscle satellite cells within the tumor stroma. These discoveries are supported by differential expression analysis and cross-dataset correlation. In the mouse embryo dataset, scMustree further uncovers spatial distribution patterns among distinct cell types, validated using spatial coordinate information. Collectively, these results establish scMustree as a powerful tool for multi-scale single-cell analysis across diverse biological contexts.

## Results

### Overview of scMustree

Single-cell RNA sequencing (scRNA-seq) data typically encompass a diverse repertoire of cell types organized in a multi-scale hierarchy, each exhibiting distinct functional traits and varying in abundance. In contrast to conventional approaches that analyze scRNA-seq data based on cell discrete partitions, scMustree is the first to treat single-cell analysis as a comprehensive exploration of tree topology, capturing both global structures and fine-grained cellular relationships. This approach naturally reveals intercellular functional relationships through branching patterns, enabling multi-scale and multi-resolution analysis of cellular composition and providing deeper insights into cellular heterogeneity and functional diversity.

The first critical step in tree construction is leaf node determination. Unlike methods such as scSHC that treat individual cells as fundamental units, scMustree implements a top-down iterative refinement strategy to generate reliable decomposed clusters (referred to as leaf nodes). Specifically, it performs cluster-adaptive dimensionality reduction followed by graph-based clustering, enabling the progressive decomposition of parent clusters through the identification of intra-cluster discriminative features. This iterative process enhances the discrimination of challenging subtypes and rare populations that might be indistinguishable through single-pass clustering, while simultaneously improving cluster purity.

Once the leaf nodes are determined, scMustree proceeds to construct the tree through a biologically informed merging strategy. Traditional methods, such as Ward’s linkage, rely on global feature comparison and often face challenges in high-dimensional spaces. To address this, scMustree first identifies differentially expressed genes (DEGs) for each decomposed cluster, which serve as functional representations of the cluster. Based on the expression patterns of these DEGs, decision trees are then constructed to rank all clusters, yielding a cluster-specific functional ranking (CFR). The CFR is a ranked permutation of all clusters derived from the decision trees trained on the DEGs of a given cluster. It reflects how other clusters are positioned relative to the functional signature of the reference cluster. Using the CFR derived from each cluster, scMustree calculates pairwise rank correlations to quantify functional similarity between clusters. These correlations guide a bottom-up merging that progressively builds the final hierarchical tree structure. This approach ensures that the resulting hierarchy reflects the functional relationships between cell types. Fig. 1 illustrates the scMustree pipeline, with detailed methodological explanations provided in the Methods section.

**Fig. 1.**
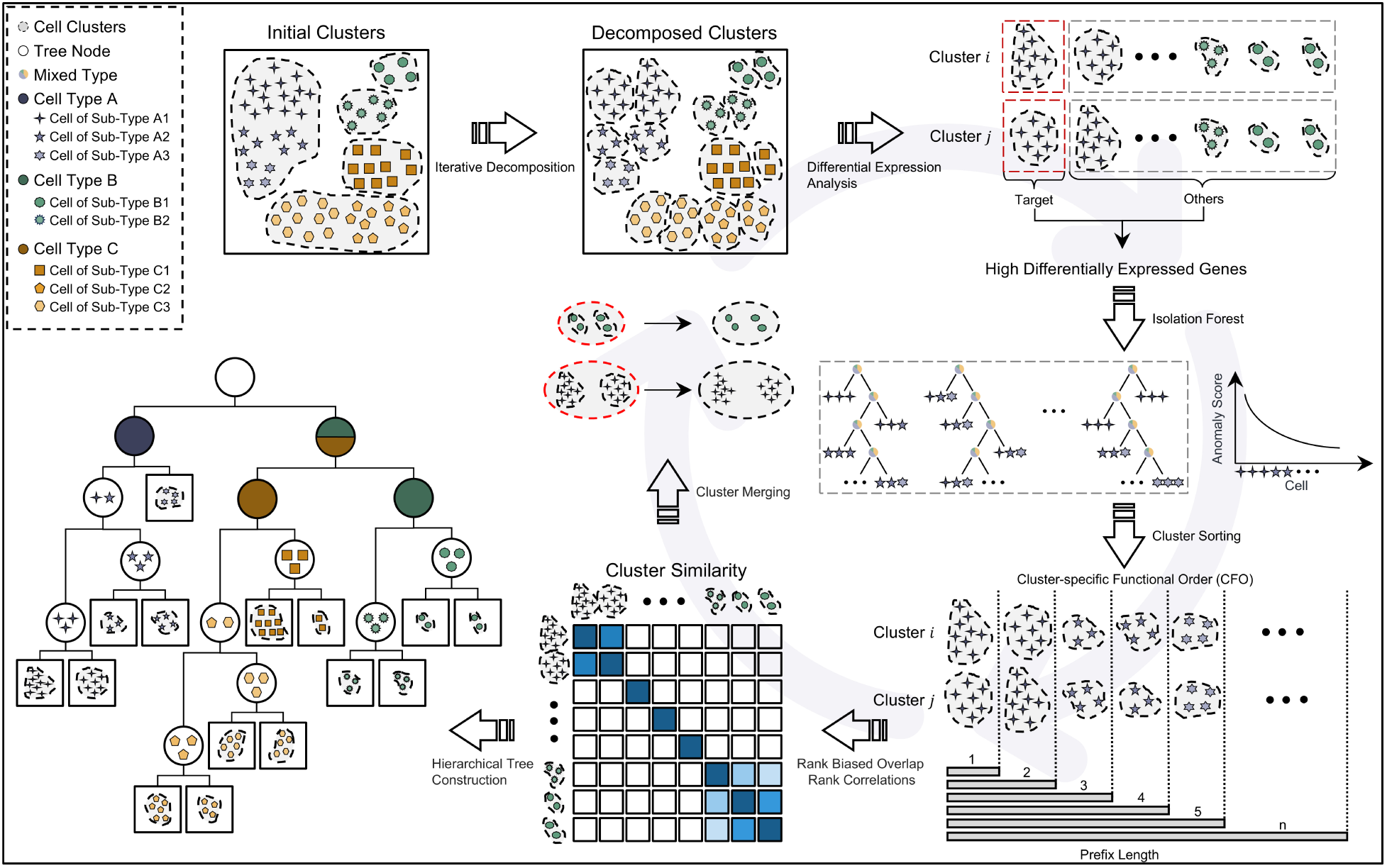
Overview of scMustree. The scMustree pipeline begins with top-down iterative clustering, decomposing initial clusters into reliable decomposed clusters (leaf nodes). Next, it identifies differentially expressed genes (DEGs) for each cluster, constructs decision trees based on their expression patterns, and defines the cluster-specific functional ranking (CFR). Using pairwise CFR rank correlations, scMustree performs bottom-up hierarchical merging, resulting in a tree structure that captures functional relationships between cell types.

### Benchmarking cellular tree construction

To comprehensively assess the performance of cellular tree construction methods, we conduct benchmarking through algorithmic accuracy and biological interpretability. For algorithmic comparisons, we select a range of representative methods, including the multi-resolution-based Louvain algorithm, dimensionality-reduction-based hierarchical clustering (scSHC), structure entropy optimization (SEAT), and deep learning-based approaches such as scDCC and scGNN. Our experiments utilize 11 single-cell datasets, covering diverse anatomical regions (e.g., airway, hippocampus) and species models (e.g., human, mouse). Notably, a subset of cell type annotations in these datasets are validated through experimental techniques such as immunofluorescence staining, flow cytometry sorting, and cross-platform dataset consistency analysis, ensuring high biological credibility. The specifics of these datasets can be found in Supplementary Table 1.

Algorithmic accuracy is quantified using normalized mutual information (NMI) and adjusted Rand index (ARI) against cell type annotations. scMustree achieves the best overall performance, with average NMI and ARI values of 0.853 and 0.861, outperforms the second-ranked method (SEAT: NMI=0.803, ARI=0.774) by 6.23% and 11.24% respectively (Fig. 2a). Additionally, we visualize the tree structures generated by the four tree-based methods across all datasets, coloring nodes according to the cell types they primarily represented. Visualization results for the hippocampus dataset are shown in Fig. 2b, with visualizations for other datasets provided in Supplementary Fig. 1. scMustree achieves branch-level type discrimination, with each major branch predominantly containing a single cell type.

**Fig. 2.**
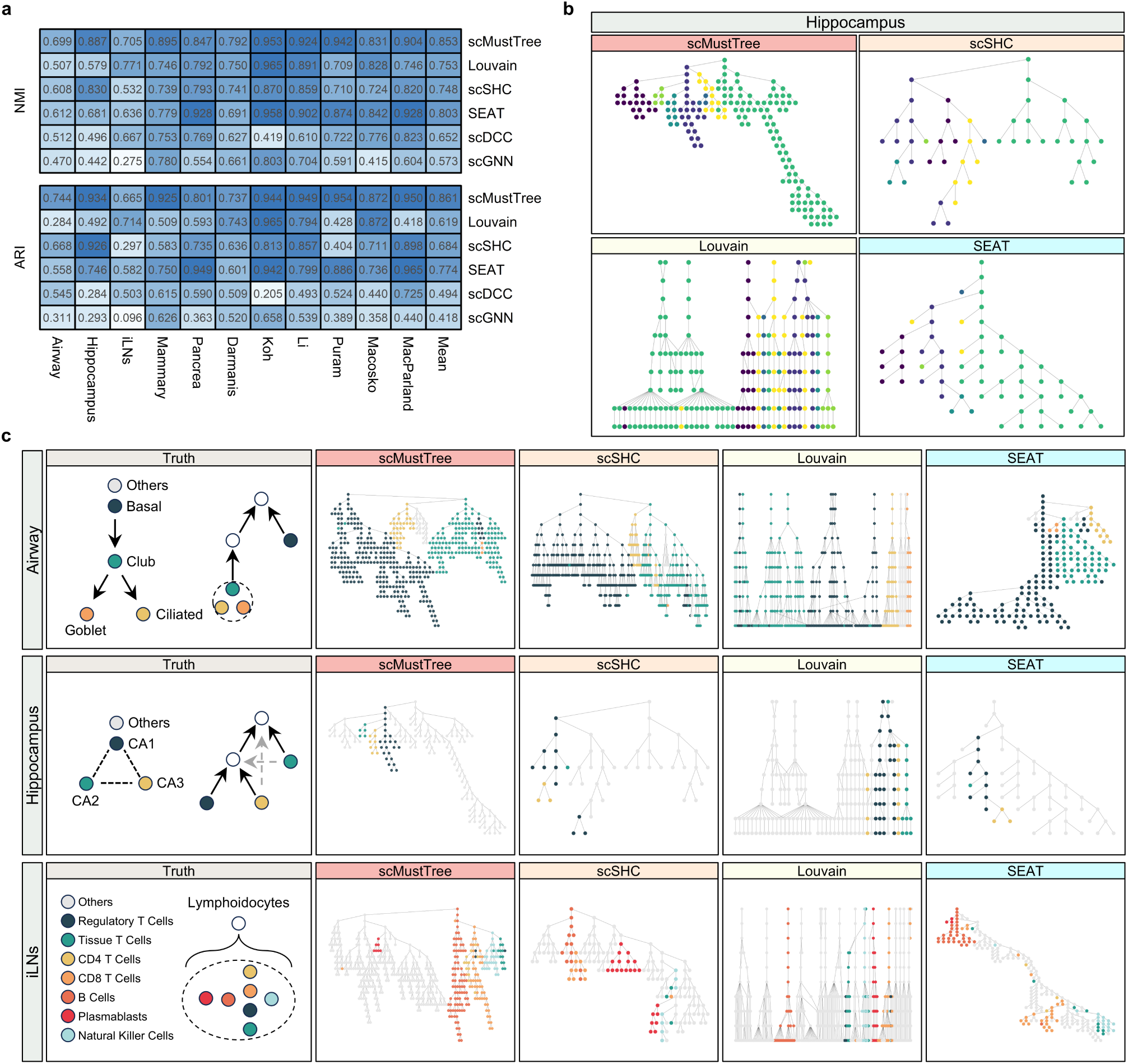
Benchmarking cellular tree construction methods in algorithmic precision and biological interpretability. **(a)** Algorithmic precision comparison of four tree-based methods (Louvain, scSHC, SEAT, scMustree) and two deep learning-based approaches (scDCC, scGNN) in terms of NMI and ARI. **(b)** Cell type-resolved tree structures generated by tree-based methods in the Hippocampus dataset. Nodes are colored by dominant cell types. **(c)** Biological interpretability assessment across three datasets with reliable ground-truth relationships (left panels) and corresponding method performance (right panels).

To evaluate the biological interpretability of constructed trees, we analyze the topological consistency between tree architectures and reliable cell-type relationships derived from original dataset studies and known biological relationships. We use three representative datasets with well-defined cellular relationships: (1) Developmental trajectories (Airway): lineage-committed transitions from basal to secretory cells; (2) Spatial subtypes (Hippocampus): Anatomically distinct neuronal subtypes (CA1∼CA3); (3) Functional modules (iLNs): Lymphocyte subtypes with coordinated immune responses.

The airway dataset published by Montoro et al. is predominantly composed of basal cells and club cells^30^. To investigate differentiation dynamics, the authors developed an innovative “pulse-seq” methodology that integrates single-cell RNA sequencing (scRNA-seq) with in vivo genetic lineage tracing, enabling temporal monitoring of cell fate transitions. Their lineage tracing analysis revealed a hierarchical differentiation pathway: basal cells initially differentiate into club cells, which subsequently generate goblet and ciliated cells (Fig. 2c). Among the tree structures generated by the four methods, the Louvain algorithm, operating through multi-resolution clustering, faces inherent constraints in lineage reconstruction. While lower clustering resolutions naturally yield fewer cell groups, this parameter reduction does not guarantee biologically meaningful merging of distinct cell types. While SEAT captures the overall differentiation hierarchy, it inaccurately positions goblet and ciliated cells of equivalent differentiation states at distant branches. scSHC exhibits structural similarities to scMustree in preserving differentiation relationships. However, scSHC incorrectly clusters other cell types primarily along the branch dominated by club cells. In the Hippocampus dataset^31^, focusing on the CA1, CA2, and CA3 subregions that form interconnected neuronal circuits, scMustree successfully clusters these subregions into three distinct branches within a unified subtree, aligning with their known biological interplay. In contrast, scSHC and Louvain disperse these subregions across unrelated branches, while SEAT clusters them within a single branch but fails to maintain CA1 spatial coherence. Analysis of the iLNs dataset^32^, which contains diverse immune cell populations, specifically highlights scMustree’s precision in segregating lymphocyte types into discrete branches of a shared subtree. Other methods exhibit erroneous adjacency relationships between distinct lymphocyte types (Fig. 2c).

In summary, scMustree demonstrates significant advantages in both algorithmic accuracy and biological interpretability. Its superior clustering performance and accurate representation of biological relationships between cell types establish scMustree as a powerful tool for constructing multi-scale cellular tree structures in single-cell data analysis.

### scMustree yields high-purity leaf nodes

Accurately determining the leaf nodes is critical for constructing a reliable tree structure. To evaluate the accuracy of the decomposed clusters (as leaf nodes) obtained by scMustree, we conduct a quantitative analysis of the accuracy of clusters before and after decomposition. For each cluster, we first calculate its purity based on the annotation information, defined as the proportion of the most abundant cell type within the cluster. The purity of cluster *i* is calculated as follows: 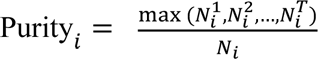, where *N* is the total number of cells in cluster *i*, *N*^*i*^ is the number of cells of type *j* in cluster *i*, and *T* is the total number of cell types in the dataset. We compute the average cluster purity distribution before and after decomposition for all datasets (Fig. 3a). The average purity for each dataset is shown in Fig. 3b. The results demonstrate that most leaf nodes are predominantly composed of a single cell type.

**Fig. 3.**
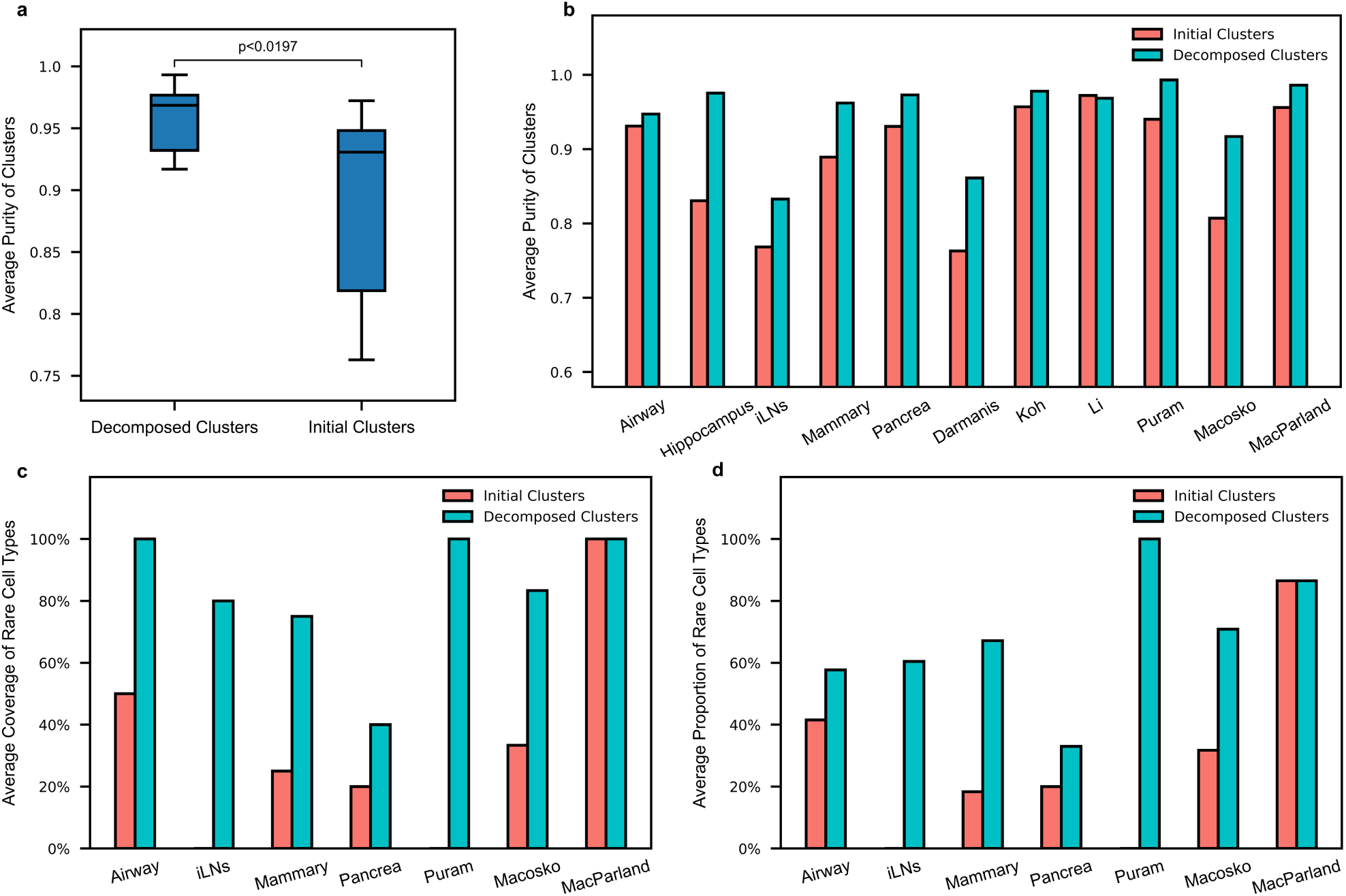
Benchmarking scMustree’s performance in leaf node determination. (a) Distribution of average cluster purity before and after decomposition. (b) Average cluster purity for each dataset before and after decomposition. (c) Average coverage of rare cell types before and after decomposition. (d) The average proportion of cells involved in rare cell types before and after decomposition.

For datasets containing rare cell types (<1% abundance), we develop a rare-type coverage metric. First, we identify the most abundant cell type in each cluster, remove duplicates, and obtain the set of types before and after decomposition, denoted as *C*. The set of rare cell types in the data is defined as *c_rate._* The coverage of rare cell types is calculated as 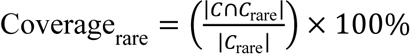. Fig. 3c reveals that decomposed clusters achieve substantially better rare-type coverage than initial clustering, with previously undetected rare types in some datasets being effectively identified.

Additionally, we examine the proportion of cells involved in clusters primarily composed of rare cell types. This proportion is calculated as follows: 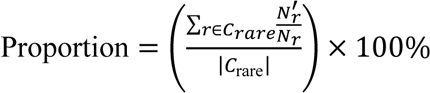, where *N*_r_ is the total number of cells of rare type *r* in clusters primarily composed of type *r*, and *N*_r_ is the total number of cells of rare type *r*. The average proportion of cells involved in rare cell types for each dataset before and after decomposition is visualized in Fig. 3d. The results indicate that most major rare cell types (with a proportion greater than 60%) are effectively separated after decomposition. In summary, scMustree achieves high-purity leaf nodes by iteratively decomposing initial clusters through hierarchical clustering.

### scMustree identifies multi-scale cell subtypes in the tumor microenvironment

Transcriptomic profiling has revolutionized our understanding of regulatory programs and disease subtypes across human malignancies^33,34^. Emerging evidence highlights that functional diversification of non-malignant cells within the tumor microenvironment (TME) critically influences tumor progression and therapeutic responses^35,36^. To investigate this cellular complexity, we analyze the Puram dataset comprising 3,363 non-malignant cells from 18 head and neck squamous cell carcinoma (HNSCC) patients^37^. Initial clustering categorizes these cells into 8 major types: T cells, B/plasma cells, macrophages, dendritic cells, mast cells, endothelial cells, fibroblasts, and myocytes. The spatial relationships of these types are visualized through UMAP dimensionality reduction (Fig. 4a).

**Fig. 4.**
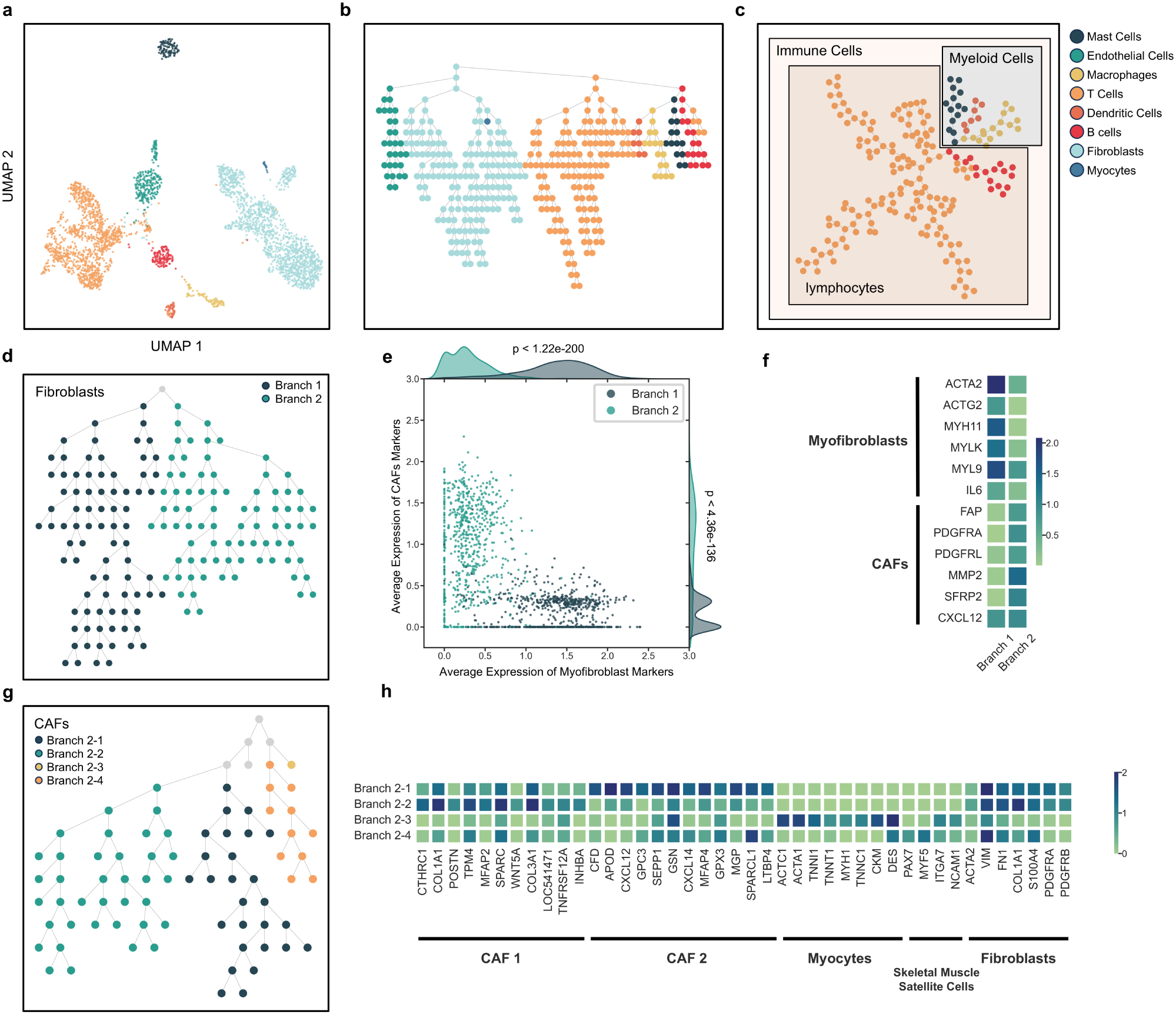
Multi-scale hierarchical analysis of tumor microenvironment (TME) cellular functional diversification using scMustree in the Puram dataset. **(a)** UMAP projection of single-cell transcriptomes color-coded by annotated cell types. **(b)** The complete tree structure is constructed by scMustree, with node colors denoting dominant cell type annotations. **(c)** Flattened representation of the immune-cell-dominated subtree, with node colors distinguishing dominant immune cell types. **(d)** Fibroblast-dominated subtree with dual-color highlighting of its two principal branches. **(e)** Joint plot showing branch-specific expression patterns of myofibroblast and CAF markers across single cells (central scatter plot) with marginal density distributions. Statistical significance is assessed using a two-sided Wilcoxon rank sum test, with p-value adjustment via Bonferroni correction based on the total number of genes. **(f)** Comparative visualization of mean marker expression levels between branches for myofibroblast and CAFs. **(g)** CAFs-dominated subtree with quad-color differentiation of its four major branches. **(h)** Comparative visualization of mean marker expression levels between branches for five cell types.

Applying scMustree to this dataset reconstructs a hierarchical tree structure that comprehensively maps functional relationships between cell types across multiple scales (Fig. 4b). This architecture aligns with established cell type annotations, as evidenced by the distinct expression patterns of markers in branch-specific nodes (Supplementary Fig. 2; marker lists in Supplementary Table 2). Within the scMustree tree, we examined the expression patterns of a specific or functionally known gene set, *G*_sub_. Specifically, for each gene *G*_i_ in *G*_sub_, the mean (μ) and standard deviation (σ) of its expression were calculated. Then, each cell’s expression value for gene *G*_i_ was standardized using a Z-score, computed as: 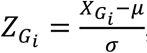 is the expression vector of gene *G_i_* across all cells. The expression score for node *D*_j_ is then calculated as: 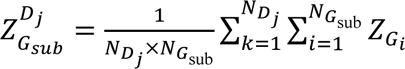, where *N*_*Di*_ is the number of cells in node *D*_*j*_ and *N*_*G*sub_ is the number of genes in the gene set. The scores are subsequently min-max normalized and divided into 10 groups for quantitative visualization using different colors.

Beyond validating established classifications, the tree topology reveals functional relationships and intra-population heterogeneity. For instance, macrophages and dendritic cells occupy neighboring branches due to shared myeloid lineage characteristics, and the fibroblast-dominated subtree exhibits spatial segregation corresponding to molecularly distinct subtypes. The differential expression levels of fibroblast markers within its branches point to the existence of functionally distinct subtypes.

To further validate the topology of this tree, we first examine the immune-cell-dominated subtree in its flattened representation (Fig. 4c). The structure separates lymphoid lineages (T cells, B/plasma cells) from myeloid lineages (macrophages, dendritic cells, mast cells), consistent with established immunological hierarchies. Given the emerging clinical significance of tumor-associated fibroblasts, subsequent analysis focuses on the fibroblast-dominated subtree (Fig. 4d). Through differential expression analysis (top 50 up-regulated genes in Supplementary Data 1), scMustree identifies two major fibroblast subtypes originally described in the dataset: myofibroblasts (Branch 1) characterized by elevated *ACTA2*, *MYH11*, and *MYL9* expression, and cancer-associated fibroblasts (CAFs, Branch 2) marked by *FAP*, *PDGFRA*, and *MMP2* expression. A scatter plot with marginal distributions visualizes the average expression levels of these subtype-specific markers across individual cells, color-coded by branch (Fig. 4e), with comparative average expression levels of each marker between branches in Fig. 4f.

To delineate finer functional subdivisions within CAFs, we perform progressive branching analysis along the CAF subtree, which reveals four distinct sub-branches (Branches 2-1 to 2-4, Fig. 4g). The average expression levels of each marker in these branches are shown in Fig. 4h. Differential expression analysis (top 50 up-regulated genes in Supplementary Data 2) identifies Branches 2-1 and 2-2 as corresponding to previously reported CAF1/CAF2 subtypes. Cells in Branch 2-3 exhibit myocyte-like features (*DES*^+^, *ACTA1*^+^) which are previously annotated as myocytes. Notably, after merging with Branch 2-4, this composite branch localizes adjacent to the fibroblast-dominated branches (Branches 2-1 and 2-2). Further characterization reveals that Branch 2-4 cells co-express fibroblast markers (*S100A4*^+^, *VIM*^+^) and muscle satellite cell markers (*PAX7*^+^), suggesting they may represent activated muscle satellite cells with increased gene expression related to extracellular matrix remodeling and fibrosis processes, representing a previously unrecognized cell population in the original study.

Building on this hierarchical tree, scMustree similarly enables multi-scale analysis of T cell heterogeneity. Through differential expression analysis and the analysis of the average expression of markers, we confirm that its four main branches match four reported subtypes: regulatory T cells, conventional *CD4*^+^ T helpers, and two cytotoxic *CD8*^+^ T cell states (activated and exhausted) (Supplementary Fig. 3 and Supplementary Data 3).

In summary, the tree structure constructed by scMustree not only captures the multi-scale type relationships obtained through multiple rounds of analysis in the original study but also uncovers novel TME-associated cell types. This demonstrates its utility for decoding functional diversity in complex environments.

### scMustree infers cell-type relationships validated across multiple species

Understanding cellular interactions within the aqueous humor outflow pathways provides critical insights into glaucoma pathogenesis and therapeutic development^38,39^. Previous single-cell RNA sequencing studies have established a foundational atlas of adult trabecular meshwork (TM) and adjacent tissues, identifying 19 histologically validated cell types^40^. This framework enables systematic cross-species comparisons in cynomolgus macaque, rhesus macaque, porcine, and murine models. Building upon this foundation, Mah et al. develop an evolutionary model for phylogenetic cell-type classification using multi-species datasets^41^. To facilitate a direct comparison with Mah et al.’s cross-species cell phylogeny structure, we focus on 11 consensus cell types (18,203 cells) retained in their evolutionary framework. The spatial relationships among these types are visualized through UMAP dimensionality reduction (Fig. 5a).

**Fig. 5.**
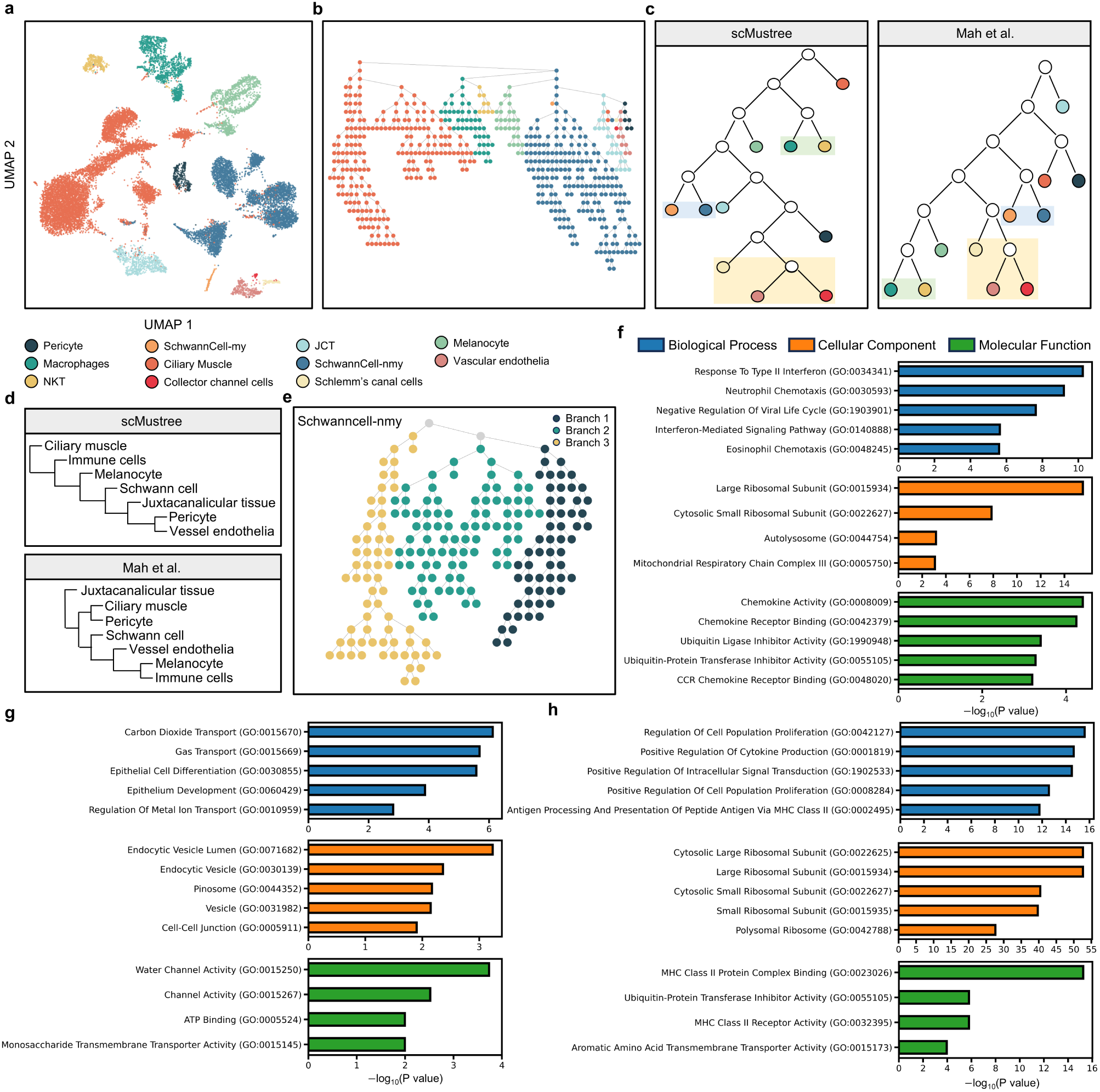
Analysis of cell type relationships and functional diversification in the human glaucoma dataset using scMustree. **(a)** UMAP projection of single-cell transcriptomes, color-coded by annotated cell types. **(b)** The complete tree structure is constructed by scMustree, with node colors denoting dominant cell type annotations. **(c)** Comparison of key branching patterns between scMustree and Mah et al.’s cross-species cell phylogeny structure in the refined topology. **(d)** Topological distinctions in inter-type relationships after integration of functionally related cell types, comparing scMustree with Mah et al.’s structure. **(e)** Non-myelinating Schwann cell-dominated subtree, with three colors highlighting its three principal branches. **(f∼h)** GO enrichment analysis results correspond to Branch 1 **(f)**, Branch 2 **(g)**, and Branch 3 **(h)**. NKT, Natural Killer T cells; my, myelinating; nmy, non-myelinating; JCT, juxtacanalicular canal tissue.

scMustree reconstructs a multi-scale hierarchical tree that fully captures the relationships among the 11 cell types (Fig. 5b). We further refine the topology to compare key branching patterns (Fig. 5c), thereby showcasing scMustree’s capacity to identify biologically meaningful proximities that align with cross-species phylogenetic frameworks. Specifically, immune cell types (macrophages and NKT cells) form adjacent branches reflecting shared immune surveillance functions^42^. Schwann cell subtypes (myelinating and non-myelinating) are grouped according to developmental lineage^43^. Meanwhile, the ‘vessel endothelia’ types (endothelia, Schlemm’s canal, collector channel cells) share canonical endothelial markers, following the classification by van Zyl et al^40^.

A notable divergence emerges in pericyte positioning. Mah et al. associate pericytes with the ciliary muscle, but scMustree positions them adjacent to vascular endothelia. This topological realignment more accurately reflects in vivo biology, as pericytes positioned adjacent to vascular endothelia regulate capillary dynamics and blood flow, aligning with their well-established role in vascular homeostasis^44^. In contrast, the ciliary muscle primarily contributes to tissue structure and contractile function essential for lens accommodation^45^, making its association with pericytes less consistent with their known physiological function.

We integrate the above functionally related cell types to reduce redundancy among functionally overlapping types, allowing for clearer observation of inter-type distinctions in topological comparisons (Fig. 5d). The hierarchical merging in scMustree starts with the integration of pericytes and vascular endothelia, which is anatomically justified by their shared basement membrane localization. It then progressively incorporates juxtacanalicular tissue, Schwann cells, melanocytes, immune clusters, and ciliary muscle. This stepwise integration pattern recapitulates the spatial gradient from the vascular core (pericytes-endothelia) to adjacent outflow structures (JCT), ultimately reaching distal contractile elements (ciliary muscle).

Detailed analysis of non-myelinating Schwann cells (SchwannCell-nmy) reveals three functionally distinct subclusters (Fig. 5e). Differential expression analysis (Supplementary Data 4) identifies branch-specific markers, coupled with Gene Ontology (GO) enrichment (Fig. 5f∼h; complete results in Supplementary Data 5∼7) confirming functional divergence. Branch 1 demonstrates stress-response specialization, highlighted by molecular chaperones *CRYAB* and *HSPB1*, interferon regulators *IFITM3* and *ISG15*, and chemokine signaling through *CCL2*. GO terms “Response to Type II Interferon” and “Ribosomal subunit” reflected inflammation-adaptation duality. Branch 2 exhibits epithelial-ionic regulation with *KRT13* and *KRT19* for barrier maintenance, *CA2/CA4* mediating “Carbon dioxide transport”, and *AQP1/3* maintaining osmotic balance. Stress-adaptation genes *HSPA1B* and *ZFP36L1* suggest microenvironmental modulation. Branch 3 specializes in immune coordination, dominated by antigen presentation genes *HLA-DRA* and *CD163*, inflammatory regulators *CEBPD* and *TSC22D3*, alongside translation machinery *RPL/RPS* family. Enriched “MHC class II antigen presentation” and “Regulation of cytokine production” confirm adaptive immune crosstalk.

These findings demonstrate scMustree’s multi-scale capability to validate evolutionarily cell-type relationships while resolving functional diversity within cellular types. The hierarchical architecture preserves the global topological features of TM ecosystems while enabling granular analysis of subtype-specific pathophysiology, providing a powerful framework for linking functional diversity to disease mechanisms.

### scMustree identifies Alzheimer’s-related microglial subtypes in the human hippocampus

The diversity in the functions of brain cell types is of great significance for gaining a deeper understanding of neurodegenerative diseases such as Alzheimer’s disease (AD)^46^. Leveraging an Alzheimer dataset comprising 118,240 cells across 13 cell types from human brain samples (hippocampus and superior/middle temporal gyrus (SMTG)) of six AD patients and six matched controls, we apply scMustree to investigate cellular dynamics^47^. The spatial relationships among cell types are visualized through UMAP dimensionality reduction (Fig. 6a).

**Fig. 6.**
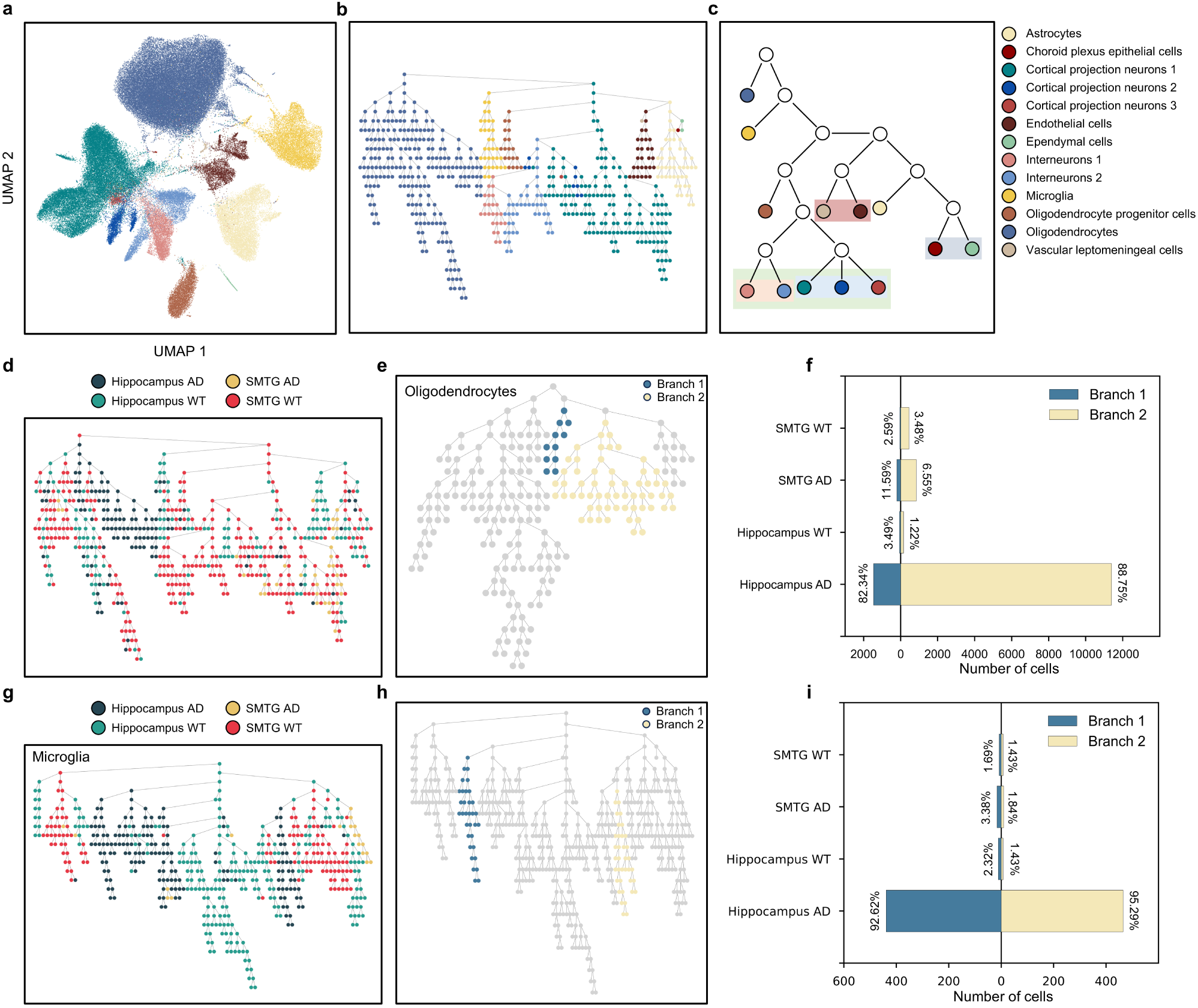
Analysis of cell type relationships and functional diversification in the Alzheimer dataset using scMustree. **(a)** UMAP projection of single-cell transcriptomes, color-coded by annotated cell types. **(b)** The complete tree structure is constructed by scMustree, with node colors denoting dominant cell type annotations. **(c)** Key branching patterns in the refined topology of scMustree’s tree. **(d)** The complete tree structure is constructed by scMustree, with node colors representing the sources of cells, including cells from Alzheimer’s patients (AD) and wild-type (WT) in the two brain regions (hippocampus and superior/middle temporal gyrus (SMTG)). **(e)** Oligodendrocyte-dominated subtree highlighting two branches predominantly composed of AD hippocampal cells. **(f)** Composition analysis of cell sources across oligodendrocyte sub-branches from AD hippocampus. **(g)** Microglia-specific tree structure constructed by scMustree with cell sources (AD/WT in hippocampus/SMTG). **(h)** Microglia subtree highlighting two AD hippocampal-enriched branches. **(i)** Comparative cell source composition in microglial sub-branches from AD hippocampus.

scMustree constructs a multi-scale hierarchical tree that captures cell type relationships (Fig. 6b), with refined topological analysis showcasing its capacity to identify adjacent branches of cell types that may reflect functional modules potentially linked to AD pathogenesis (Fig. 6c).

Specifically, three cortical projection neuron subsets (1, 2, and 3), and two interneuron subsets (1 and 2) are adjacent after each being merged into their respective main branches. Cortical projection neurons are responsible for long-distance signal transmission, while interneurons regulate local inhibitory circuits. Together, they maintain the balance of neural networks. The adjacency reflects their complementary functional roles in the neural network. Vascular leptomeningeal cells and endothelial cells are both involved in the vascular system, especially the function of the blood-brain barrier. In AD, vascular lesions and blood-brain barrier leakage are typical pathological features^48,49^. Therefore, their adjacency may imply the synergistic effect between these cell types, affecting the pathological process of AD. Choroid plexus epithelial cells and ependymal cells jointly participate in the circulation of cerebrospinal fluid and the clearance of metabolic waste. In AD, the accumulation of beta-amyloid protein is closely related to the impaired function of cerebrospinal fluid circulation and waste clearance^50^.

Furthermore, we observe the distribution of cells from Alzheimer’s patients (AD) and wild-type (WT) in the two brain regions throughout the entire tree (Fig. 6d). Interestingly, we find two main branches of oligodendrocytes composed of cells in the hippocampus of AD patients, Branch 1 and Branch 2, accounting for 82.34% and 88.75% respectively (Fig. 6e and f). To explore their potential relevance to AD, we conduct differential gene expression analysis and GO enrichment analysis to analyze their functions (Supplementary Fig. 4a and b, complete results in Supplementary Data 8∼10), which show distinct functional profiles, highlighting their potential roles in the disease. Branch 1 is mainly characterized by the high expression of heat shock protein family genes (such as *HSP90AA1*, *HSPH1*, *HSPA1A*) and the lysosomal sorting receptor *SORL1*, which is significantly enriched in the negative regulation pathways of the unfolded protein response (UPR) and inclusion body assembly. This expression pattern may resist the Aβ production and tau phosphorylation pathology in the early stages of AD by stabilizing tau protein kinase activity (inhibiting the over-activation of GSK-3β) and reducing the abnormal processing of amyloid precursor protein (*APP*). Branch 2 is centered on metabolic-inflammatory regulation. Its highly expressed genes such as *ACSL1*, *HIF1A*, and *SPP1* drive hypoxia-induced glycolytic reprogramming (such as the up-regulation of *PFKFB3*) and pro-inflammatory signal transduction (such as the activation of the osteopontin *SPP1*-integrin-*ERK* pathway). While compensatorily clearing metabolic waste, these mechanisms may amplify neurodegenerative changes (such as the over-activation of microglia) and blood-brain barrier damage (*HIF1A-VEGF* axis) through chronic inflammation.

Regarding microglia, which also play a complex and multifaceted role in AD, we use scMustree to reconstruct the tree structure of microglia (Fig. 6g). Combining the distribution of cells, we also find two main branches of microglia composed of cells in the hippocampus of AD patients, Branch 1 and Branch 2, accounting for 92.62% and 95.29% respectively (Fig. 6h and i). To explore their potential relevance to AD, we conduct differential gene expression analysis and GO enrichment analysis to analyze their functions (Supplementary Fig. 4c and d, complete results in Supplementary Data 11∼13), which show distinct functional profiles, highlighting their potential roles in the disease. Interestingly, we find that Branch 1 of microglia and the previously discovered Branch 1 of Oligodendrocytes show very similar differential genes and functions, which indicates that different glial cell types in the hippocampus of AD patients may work together to cope with pathological stress through conserved stress-response mechanisms. As for Branch 2, it shows characteristics closely related to metabolism and inflammation. Its highly expressed genes such as *ACSL1*, *HIF1A*, and *SPP1* drive hypoxia-induced glycolytic reprogramming and the activation of pro-inflammatory signals. This metabolic reprogramming may lead to the accumulation of lipid droplets, which is closely related to intracellular lipid metabolic disorders. Therefore, the microglia of Branch 2 may promote the accumulation of lipid droplets through these metabolic changes, thereby affecting cell function and the pathological process of AD. Interestingly, after research, we find that this subtype is very similar to the lipid-associated *ACSL1*^+^ microglia subtype reported by Haney et al.^51^ They report LD-accumulating microglia (LDAM), a state of disease-associated microglia (DAM) microglia, which co-expressing of more metabolic state regulators such as *NAMPT* and *DPYD*. We visualize the expression distribution of LDAM and DAM-related markers in the tree structure, further confirming their identities (Supplementary Fig. 5).

In conclusion, scMustree can not only comprehensively reflect the relationships among cell types in disease states but also display disease-related cell type subtypes, which play different functions in the disease, thus providing further insights for disease analysis.

### scMustree effectively characterizes dendritic and bipolar cell subtypes through tree construction

In cellular biology research, the precise identification and characterization of cellular subtypes has become paramount for understanding functional diversity. This is particularly crucial for dendritic cells (DCs) and bipolar cells (BCs), where subtype variations determine their distinct roles in immune responses^52^ and neural signaling^53^, respectively. Building upon previous subtype characterization efforts, we demonstrate how scMustree’s hierarchical tree architecture effectively resolves cellular subtypes in these critical cell subtypes.

Using 2,099 DCs extracted from the 68k PBMC dataset, we evaluate scMustree’s ability to identify subtypes beyond original annotations. UMAP visualization reveals spatial relationships among DCs (Fig. 7a), while scMustree generates a multi-scale hierarchical tree with five principal branches (Fig. 7b). To validate these branches, we perform cross-dataset correlation analysis comparing the transcriptional profiles of cells within each branch against Villani et al.’s FACS-validated DC subtypes^54^ using their established marker set (Fig. 7c, marker list in Supplementary Table 3). The analysis reveals distinct subtype associations: Branches 1 and 2 exhibit the strongest correlations with DC1 (r=0.778) and DC2 (r=0.720) respectively, while Branches 3 and 4 both show high similarity to DC4 (r=0.836 and 0.754). Notably, Branch 5 demonstrates significant alignment with plasmacytoid DCs (pDCs/DC6, r=0.777). The observed transcriptional gradients between adjacent branches suggest a topological hierarchy in subtype relationships, potentially reflecting developmental or functional continuities within the DC subtype.

**Fig. 7.**
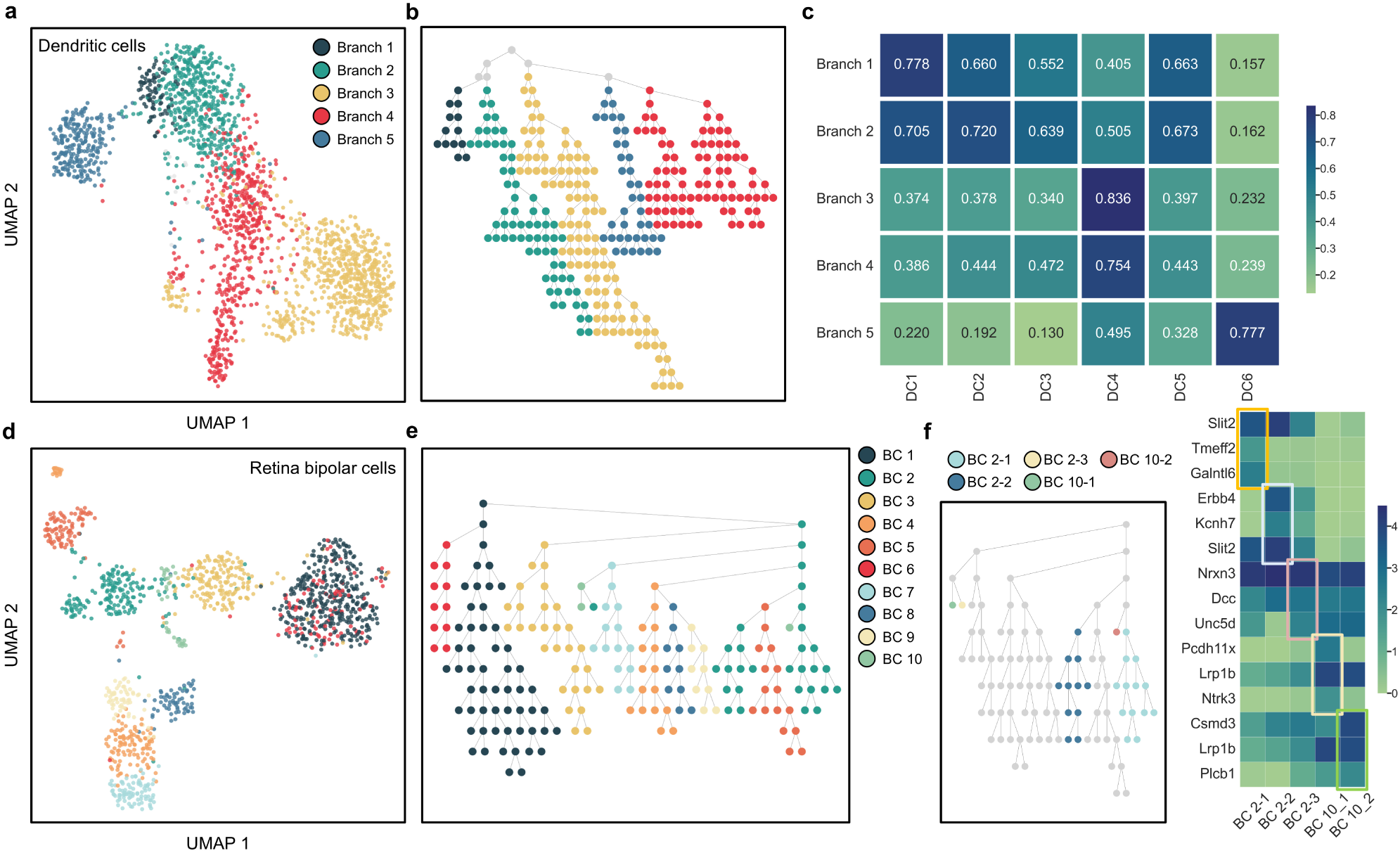
Analysis of dendritic cells (DCs) in human PBMCs and bipolar cells (BCs) in mouse retinal datasets using scMustree. (a) UMAP projection of single-cell transcriptomes, with coloring based on different branches of the scMustree tree structure. (b) The complete tree structure of DCs constructed by scMustree, with node colors denoting major branches. (c) Cross-dataset validation showing the similarity of marker expression patterns between established DC subtypes and corresponding scMustree-identified branches. (d) UMAP projection of single-cell transcriptomes, color-coded by BC subtypes. (e) The complete tree structure of BCs constructed by scMustree, with node colors denoting dominant cell subtype annotations. (f) Heatmap visualization of differentially expressed genes across scMustree-identified sub-branches.

Analysis of 6,198 BCs from the mouse retina dataset, which were originally annotated as ten subtypes^55^, reveals scMustree’s capacity to not only recapitulate known subtypes but also identify novel subdivisions. UMAP visualization reveals spatial relationships among BCs (Fig. 7d), while scMustree generates a multi-scale hierarchical tree with clear branch correspondence to annotated subtypes (Fig. 7e). Additionally, we observe heterogeneous subtypes, like BC2 splitting into three branches and BC10 into two. Differential gene expression analysis is done on each branch (Fig. 7f, Supplementary Data 14). Results show that though branches of the same subtype share certain gene expression patterns, they also have remarkable gene expression differences, suggesting they may have different functional roles.

These results position scMustree as a powerful tool for multi-scale subtype discovery, capable of both validating established classifications and revealing novel cellular subdivisions through its hierarchical analytical framework.

### scMustree reveals spatial distribution relationships of cell types

The spatial distribution patterns of cell types are closely associated with their physiological functions. To validate scMustree’s capability in deciphering spatial distribution relationships between cell types, we apply it to the sci-Space dataset^56^ containing 14 sagittal sections from two E14.0 mouse embryos, which provides spatial coordinates and whole transcriptome data for approximately 120,000 nuclei. Focusing on Slide 14 with the highest number of recovered nuclei (∼17,000), scMustree constructs a hierarchical tree based solely on RNA modality data without spatial coordinate reference. Using the mouse embryo anatomical regions as a reference, we analyze the spatial distribution of cells within tree nodes (Fig. 8a).

**Fig. 8.**
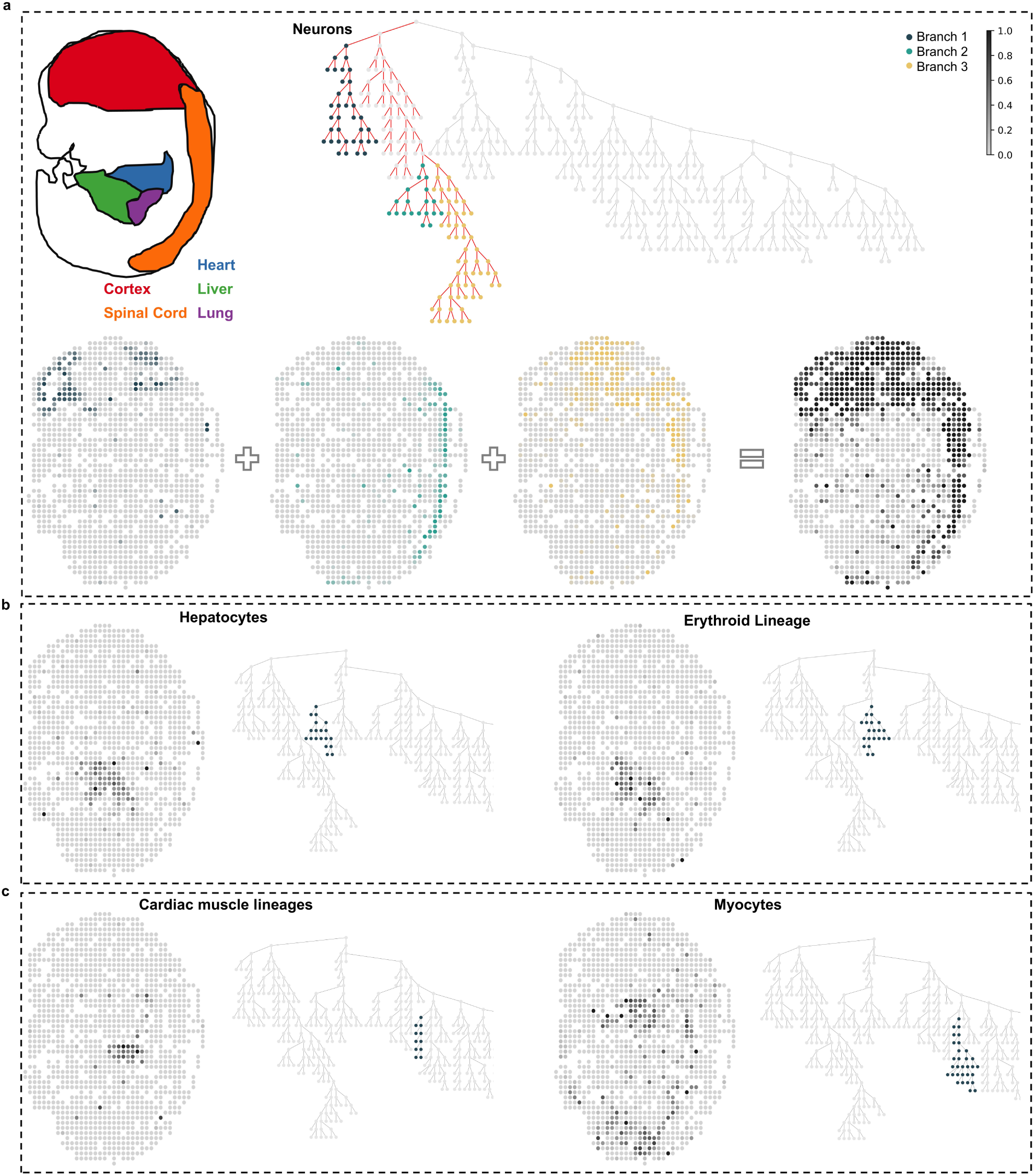
Analysis of spatial distribution and functional associations of cell types revealed by scMustree. **(a)** Anatomical regions of Slide 14 in mouse embryo sagittal sections (left) and the tree structure generated by scMustree (right). The neuron-dominated subtree is highlighted with red edges, and three main branches are color-coded. The spatial distribution of cells in the corresponding branches and subtree is shown at the bottom. Spots are marked with a continuous gradient of the same color to indicate the proportion of cells from the same branches or subtree at each spatial coordinate. **(b)** Spatial distribution patterns of cells within branches dominated by Hepatocytes and Erythroid Lineage cells. **(c)** Spatial distribution patterns of cells within branches dominated by cardiac muscle lineages and myocytes.

Specifically, we quantify the proportion of cells in each spatial coordinate within the nodes of the tree’s subtrees and visualize their spatial distribution using a continuous color gradient. Our analysis reveals a subtree in which nodes are predominantly distributed in the Cortex and Spinal Cord, and these nodes are composed of neurons. Further examination of the spatial distribution of cells within the main branches of this subtree (Fig. 8a(bottom)) shows significant spatial associations between neighboring branches. Similarly, we observe another subtree where cells within nodes are primarily distributed in the liver (Fig. 8b). This subtree contains two branches composed mainly of Hepatocytes and Erythroid Lineage cells, reflecting their spatial proximity. During mouse embryo development, these cells are predominantly located in the fetal liver.

Additionally, we investigate cardiac muscle lineages and myocytes, both derived from mesodermal myogenic progenitors during embryonic development. These cell types share key transcription factors and structural proteins. Although they do not exhibit significant spatial associations, their functional relationship is reflected in the tree structure as location proximity (Fig 8c). In summary, scMustree not only reveals spatial distribution relationships among cell types but also maintains functional associations between them.

## Discussion

The introduction of single-cell RNA sequencing (scRNA-seq) technology has transformed the study of cellular heterogeneity and functional diversity within complex tissues. Traditional clustering methods, while widely used, have limitations that restrict a comprehensive understanding of cellular relationships and functions. In this study, we present scMustree, a computational framework aiming to overcome these limitations by inferring multi-scale cell trees that capture both global and local heterogeneity.

scMustree advances single-cell analysis by treating clustering as a dynamic tree discovery process rather than static partitioning. Through iterative cluster decomposition and biologically informed merging strategies, scMustree generates high-purity cell clusters and establishes functional relationships between cell types. By identifying differentially expressed genes (DEGs) for each cluster and constructing ensemble decision trees based on their expression patterns, scMustree defines cluster-specific functional ranking (CFR). These CFR guide hierarchical merging and offer insights into the expression dynamics of cluster-specific genes.

Benchmarking against a range of representative methods across diverse single-cell datasets, scMustree demonstrates superior performance in both algorithmic accuracy and biological interpretability. The framework’s ability to achieve branch-level type discrimination and accurately represent biological relationships between cell types establishes it as a powerful tool for constructing multi-scale cellular tree structures. Furthermore, scMustree’s application to complex biological systems, such as the tumor microenvironment, glaucoma, and Alzheimer’s disease, showcases its utility in uncovering novel cell types and resolving functional diversity within cellular populations.

The multi-scale hierarchical tree structures generated by scMustree not only reflect established cell type annotations but also reveal intricate functional relationships and intra-population heterogeneity. For instance, in the tumor microenvironment, scMustree identifies distinct fibroblast subtypes and their functional associations, shedding new light on the cellular complexity and its impact on tumor progression and therapeutic responses. Similarly, in the context of glaucoma, scMustree successfully reconstructs cell type relationships that align with cross-species phylogenetic frameworks, while also resolving functional diversity within specific cell types such as non-myelinating Schwann cells.

The successful application of scMustree to various diseases highlights its potential as a valuable tool for biomedical research. By enabling a deeper understanding of cellular heterogeneity and functional diversity, scMustree can facilitate the identification of disease-specific cell types and their associated molecular mechanisms, potentially guiding the development of targeted therapies and improved diagnostic strategies. Moreover, the framework’s ability to integrate multi-scale and multi-resolution analyses makes it applicable to a wide range of biological systems, from developmental biology to immunology and neuroscience.

In conclusion, scMustree offers a robust and comprehensive approach for decoding functional diversity within complex tissues. Its innovative tree modeling framework addresses the limitations of existing clustering methods and provides researchers with a powerful tool to explore the intricate cellular landscapes underlying various biological processes and diseases. Future work could further refine the algorithm to handle larger and more complex datasets, and explore its applications in emerging areas such as spatial transcriptomics and multi-omics integration.

## Methods

### Data preprocessing

We preprocess the gene expression matrix as follows. First, we filter out genes with low expression rates that may not provide effective information. Specifically, genes that are expressed in at least three cells are retained for downstream analysis. Each scRNA-seq dataset is normalized by using the log-normalization procedure including the calculation of cell-specific size factor based on the sequencing depths, and normalization. The normalized matrix is then log2- transformed after adding 1 as a pseudo-count.

### Rapid clustering module for single-cell analysis

scMustree utilizes this rapid single-cell clustering module multiple times to decompose clusters into high-purity sub-clusters (referred to as leaf nodes). Similar to previous works, scMustree first applies principal component analysis (PCA) to obtain the top principal components (PCs) that capture the most differential signals in the data. Next, an undirected cell graph is constructed using Euclidean distance and the k-nearest neighbors (KNN) algorithm. Finally, a graph-based community detection algorithm (e.g., Louvain) is employed to assign cluster labels to the cells.

### The procedure of scMustree method

scMustree integrates three core components: 1) cluster decomposition-based leaf node determination, 2) decision tree-based cluster-specific functional orders (CFR), and 3) rank correlation-guided merging. Specifically, after data preprocessing, scMustree involves the following detailed procedures.

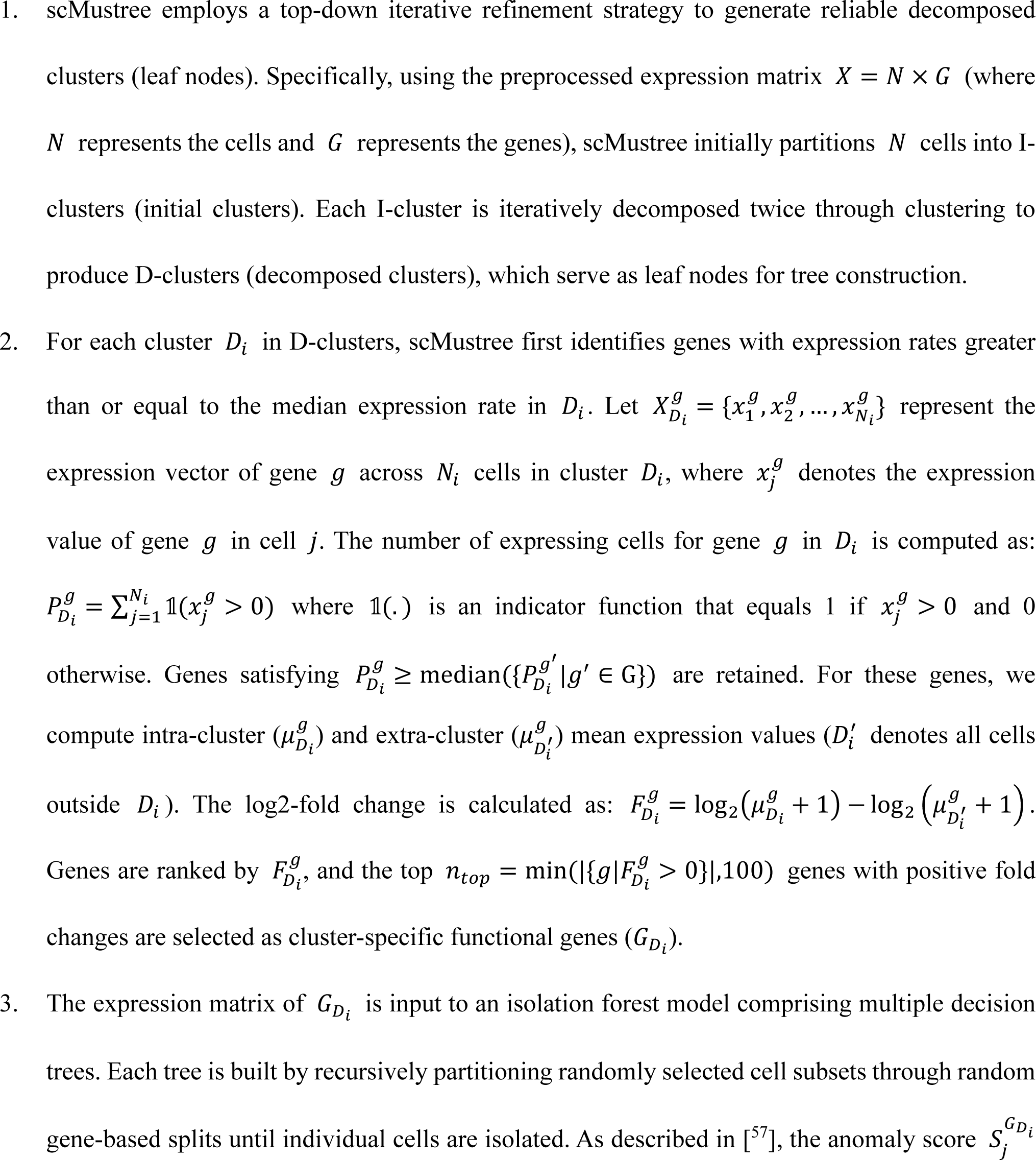
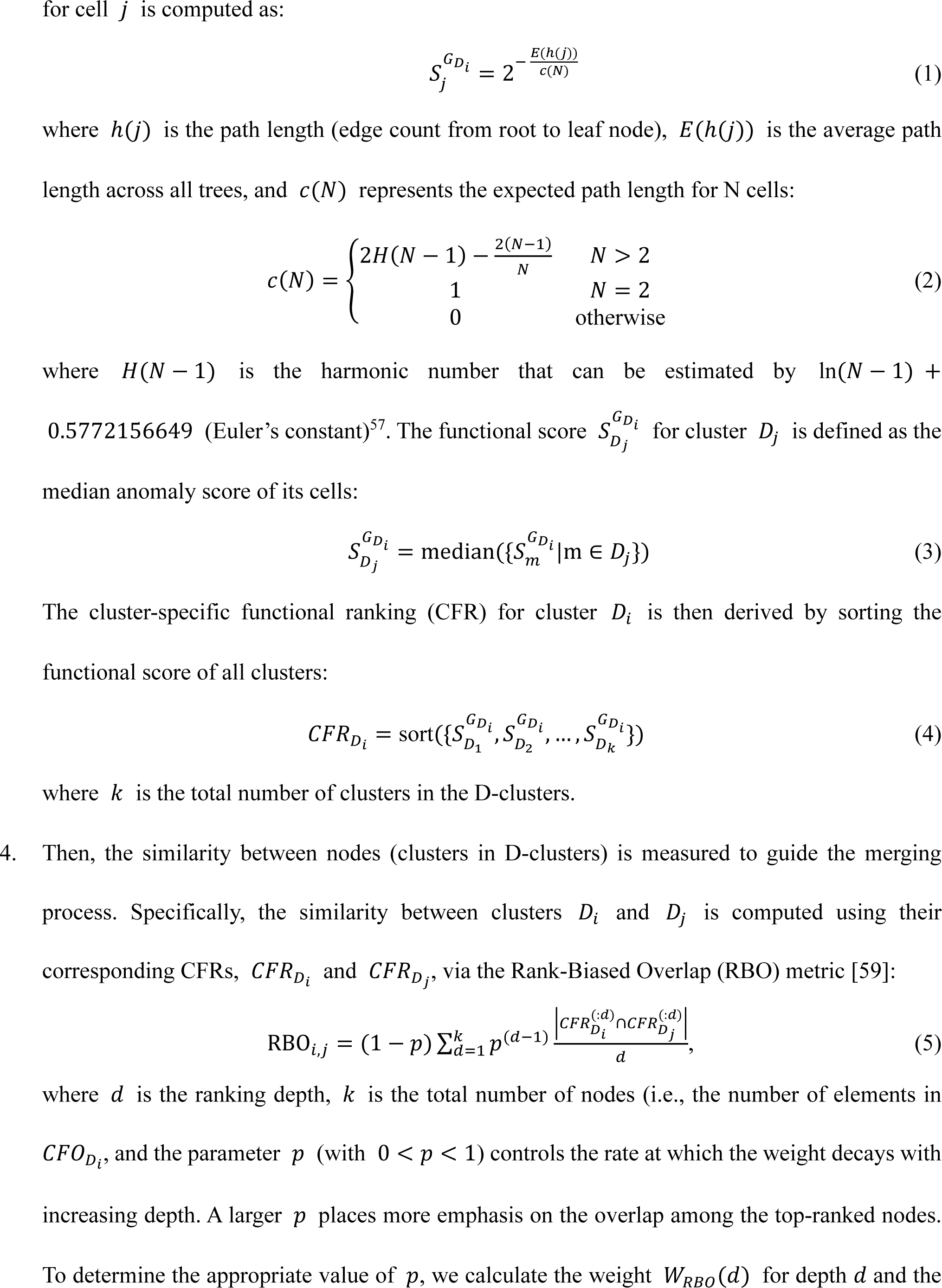
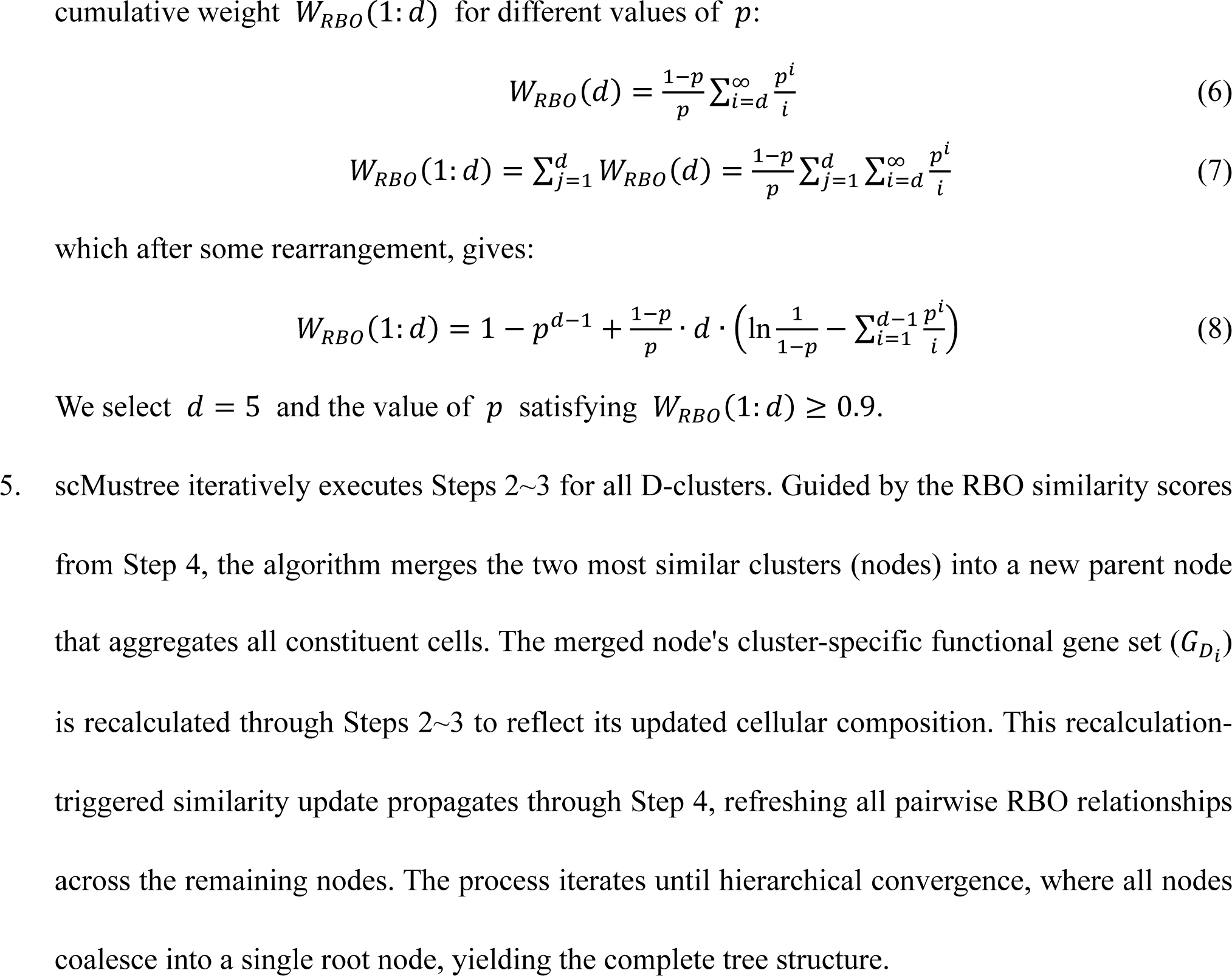

### Usage of comparative methods

To comprehensively assess the performance of cellular tree construction methods, we conduct a benchmark analysis comparing scMustree with other methods.

The multi-resolution tree structure using the Louvain algorithm is constructed by first applying the standard normalization analysis workflow from the Scanpy Python package (version 1.10.2), combined with the *best_partition* function from the community package with different resolution parameters (ranging from 0.1 to 1.0) to obtain clustering results at various resolutions. The tree is then constructed based on the overlap of clusters between adjacent resolution levels; an edge is drawn between clusters if the overlap exceeds 30%. The scSHC package is obtained from GitHub (igrabski/sc-SHC). The tree structure constructed by scSHC is generated using its internal functions and default parameters to obtain a cell distance matrix, which is computed as the Euclidean distance on the latent variables estimated by the approximate generalized linear model principal components analysis (GLM-PCA) procedure using 2,500 genes. Hierarchical clustering using Ward’s criterion is then applied to construct the tree structure. The SEAT package is obtained from GitHub (deepomicslab/SEAT). Specifically, the UMAP embedding is first obtained using the standard normalization workflow from the Scanpy Python package, and then the default parameters of SEAT are used to generate results from which the ‘Newick’ tree structure is extracted. The scDCC and scGNN packages are obtained from GitHub (ttgump/scDCC and juexinwang/scGNN, respectively). Both methods use clustering results obtained with default parameters. For some smaller datasets, we reduced the ‘pretrain_epochs’ parameter in scDCC to avoid errors, and for scDCC, the true number of cell types was provided as input.

### Identification of differential genes

A traditional Wilcoxon rank-sum test was used to identify differentially expressed (DE) genes with an FDR cutoff of 0.05 and an inter-group absolute fold-change cutoff of 1.5. Fold-change values were measured based on the group-wise mean expression values of each gene. A gene was considered cluster-specific if it was found to be differentially up- regulated in a particular cluster compared to each of the remaining clusters.

### Gene Ontology functional enrichment analysis

GO functional enrichment analysis was performed using the *enrichr* function from the gseapy Python package (version 1.1.4). The gene sets used included ‘GO_Biological_Process_2023’, ‘GO_Cellular_Component_2023’, and ‘GO_Molecular_Function_2023’.

### Statistics & Reproducibility

Differentially expressed genes were identified using the *rank_genes_groups* function from the Scanpy Python package (version 1.10.2). This analysis employed a two-sided Wilcoxon rank-sum test, with a false discovery rate (FDR) cutoff of 0.05 and an inter-group absolute fold-change cutoff of 1.5. Fold-change values were calculated based on the mean expression levels of each gene between groups. P-values are adjusted using Bonferroni correction, accounting for the total number of genes in the dataset.

No statistical method was used to pre-determine sample size. No data were excluded from the analysis, all genes in datasets were used throughout all analyses. The investigators were blinded to allocation during experiments and outcome assessment.

## Data availability

The details of the datasets used in this study are reported in Supplementary Table 1. All described datasets are obtained from various public websites under accession codes provided in Supplementary Table 1, including NCBI Gene Expression Omnibus (GEO) [https://www.ncbi.nlm.nih.gov/geo/] and Single Cell Portal [https://singlecell.zendesk.com/hc/en-us]. The Dendritic cell dataset from the 68k PBMC dataset is obtained from the website of 10X genomics ([https://www.10xgenomics.com/datasets/fresh-68-k-pbm-cs-donor-a-1-standard-1-1-0]). The Bipolar cell dataset from the mouse retina dataset is available on GitHub [https://github.com/OSU-BMBL/marsgt/tree/main/Data]. Source data are provided in this paper.

## Code availability

scMustree is publicly available at GitHub [https://github.com/xuyp-csu/scMustree].

## Acknowledgments

This work was supported in part by the National Key Research and Development Program of China (No.2021YFF1201200), the National Natural Science Foundation of China under Grants (Nos. 62350004, 62332020). This work was carried out in part using computing resources at the High-Performance Computing Center of Central South University.

## Author contributions

Y.P.X and H.D.L. conceived and designed the project. Y.P.X., S.K.W., and H.D.L. were responsible for the overall conception, design, and implementation of the study. Y.P.X. collected the datasets and conducted the experiments. Y.P.X., L.Q.D., and S.K.W. performed the data analysis. Y.P.X. tested the software operation. Y.P.X. and S.K.W. drafted the manuscript. All authors reviewed and approved the final version of the paper.

## Competing interests

The authors declare no competing interests.

## Notes

### Competing Interest Statement

The authors have declared no competing interest.

